# Effect of Electromagnetic Field (EMF) and Electric Field (EF) on Some Behavior of Honeybees (*Apis mellifera* L.)

**DOI:** 10.1101/608182

**Authors:** Yaşar Erdoğan, Mahir Murat Cengiz

## Abstract

Geomagnetic field can be used by different magnetoreception mechanisms, for navigation and orientation by honeybees. The present study analyzed the effects of magnetic field on honeybees. This study was carried out in 2017 at the Bayburt University Beekeeping Application Station. In this study, the effect of Electro Magnetic field (EMF) and electric field (EF) on the time of finding the source of food of honeybees and the time of staying there were determined. The honeybees behaviors were analyzed in the presence of external magnetic fields generated by Helmholtz coils equipment. The Electro Magnetic field values of the coils were fixed to 0 μT (90mV/m), 50 μT (118 mV/m), 100 μT (151 mV/m), 150 μT (211 mV/m), 200 μT (264 mV/m). Petri dishes filled with sugar syrup were placed in the center of the coils. According to the study, honeybees visited at most U1 (mean =21.0±17.89 bees) and at least U5 (mean =10.82±11.77 bees). Honeybees waited for the longest time in U1 (mean =35.27±6.97 seconds) and at least in U5 (mean =12.28±5.58 seconds). According to the results obtained from this first study showed that honeybees are highly affected by electromagnetic radiation and electric field.

**Summary:** Honeybees uses the magnetic field of the earth to to determine their direction. Nowadays, the rapid spread of electrical devices and mobile towers leads to an increase in man-made EMF. This causes honeybees to lose their orientation and thus lose their hives.

## INTRODUCTION

Living things have been adapted to the magnetic field of the Earth they were exposed to as a result of millions of years of natural selection. Many organisms use the magnetic field of the earth in space and time orientation (Wiltschko and Wiltschko, 2005). There is a scientific discipline called Magnetobiology, which investigates the effects of magnetic fields on living things.

Magnetobiology was formed by the unification of many scientific principles around biophysics. Magnetic fields are important ecological factors that can affect living things (Binhi and Savin, 2003; Rosen, 2003). Many studies have examined the possible effects of EMF and EF on animals. Magnetic fields and electric fields can have an impact on the daily activities, behaviors and spatial orientations of living things (Klotz and Jander, 2003; Vacha, 2006; Vacha et al., 2008). Studies have shown that ELF and EMF cause some physiological and behavioral changes on insects and increase stress protein levels. (Wyszkowska et al.2006; Wyszkowska et al. 2016). In another study carried out by Rooder (1999), they found that there was a significant increase in the motor activity of insects as EMF increased octopamine levels in insects. Honeybees are one of the most important insect affected by the electromagnetic field.

Many studies have been conducted on the response of honeybees to the electromagnetic field. They can be summarized as follows: When extra magnetic field was applied, comb building behavior and hive orientation were changed (Collett and Baron, 1994; Frier et al.1996). Free-flying honey bees can detect weak magnetic field fluctuations as much as 26 nT. It has emerged in T-labyrinth experiments where honey bees are affected by short magnetic pulses (Kirschvink, and Kobayashi-Kirschvink,1991). The magnetic remanence was detected in the abdomen of honeybees (Gould et al. 1978). Iron granules (IGs), 0.560.1 mm diameter were found in tropocytes surrounding the abdomen (Hsu and Li, 1993). Four IGs trophocyte super paramagnetic magnetite was detected under high resolution transmission electron microscopy (Hsu and Li, 1999). Fe_3_O_4_ and FeOOH were found in the honey bees (El-Jaick et al. 2001).

These results suggest that in addition to behavioral evidence, honeybees have biomagnetites necessary for magnetoreception. That is to say, honeybees have the capacity of magnetoreceptics. However, no evidence has been found to explain this capacity so far (Hsu et al. 2007).

Honeybees are of great importance for humanity and nature for many reasons. Honeybees make a great contribution to nature by providing pollination of plants other than producing beekeeping products. Therefore, bees are important pollinators for both natural vegetation and for crops (Castro, 2001). According to the studies conducted, the economic value of honeybees was about 153 billion euros in 2005 (Gallai et al., 2009). The European honeybees (*Apis mellifera* L.) is one of the most effective pollinator insects (Garibaldi et al., 2011, Garibaldi et al., 2014). In addition, *Apis mellifera* L. is the most widely used bees in the world for beekeeping.

Honeybees carry honey, pollen, propolis, and water from the outside to their hives. Honeybees are talented insects who can find plants in the field and return to the hive. Worker honeybees are rare social insects that collect foods from distances of up to 8-12 km and return to their hives without losing direction.

Honeybees use the position of the sun (Rossel and Wehner, 1984), polarized light (Rossel and Wehner, 1986; Evangelista et al., 2014) and landmarks (Dyer and Gould, 1981) to determine their direction.

The ability of the bees to feel the Electromagnetic field of the Earth is one of the most important factors that honeybees use in finding direction. Although it is thought that the most important factor that honeybees use in finding direction is the sun; they can also use **cues** such as smell, polarized light, compass of the sky, signs around the hive, chemicals, acoustic instruments and magnetic field.

The state of the sky (cloudy sky or clear blue sky) and the time of day determine which of these elements will be used by honeybees. Today, the use of devices that produce the Electro Magnetic field such as mobile towers, mobile phones, Wi-Fi, Bluetooth, electric appliances and high voltage lines has increased considerably.

The increase in these devices has led to debates that the ability of the honeybee to navigate has disappeared when the magnetic field causes negativity on human and other living things. Depending on the intensity of the magnetic field and the duration of exposure, the risk of developing cancer (Wertheimer and Leeper 1979) leukemia (Greenland et al., 2000; Draper et al., 2005), lymphoblastic leukaemia (Hatch et al., 1998), acute lymphblastic leukaemia (Kabuto et al. 2006) and alzheimer’s (Huss et al., 2006) are increased.

According to the results of the studies, it was found that humans (Leszezynski et al., 2002; Gandi and Singh, 2005; Hardell and Sage, 2008), rabbits, rats (Moorhouse and Macdonald, 2005), bats (Nicholls and Racey, 2007; Nicholis and Paul, 2007), birds (Everaert and Bauwens, 2007;Balmori, 2009; Grigoriev, 2003), frogs (Balmori, 2006; Balmori, 2010), nematodes, Drosophila (Ghamdi, 2012), plants (Belyavskaya, 2001; Haggerty, 2010), Paper wasp (Pereira-Bomfim et al. 2015), and honey bees (Harst et al., 2006; Sharma and Kumar, 2010; Favre, 2011) have been reported to be influenced by electromagnetic fields (EMF).

Pereira-Bomfim et al. (2015) showed that the social wasp *Polybia paulista* is sensitive to modifications in the local geomagnetic field. This study, which was made with magnets and Helmholtz coils equipment, showed that the change of the magnetic field affects the flight activity of *Polybia paulista* (Ihering).

Recently there have been reports of many factors affecting the development of honeybees, such as disease, natural enemies, pesticides and adverse climatic conditions (Favre, 2011).

The increase in losses in bee colonies all over the world has caused a phenomenon in which the number of bees in the hive decrease very rapidly, without showing the symptom of an illness. Scientists believe that these phenomena, called the Colony Collapse Disorder (CCD) (Gallai et al., 2009), are caused by viruses, unscientific farm applications, monoculture, no hygienic farming conditions, sudden changes in the climate, pesticides, air pollution, and even GMO crops.

At present, it is rgued that the most important cause of CCD is electromagnetic pollution (Kumar, 2018; Taye et al.,2017; Cammaerts, 2017). Due to increased electromagnetic pollution, it is suggested that the honeybees come out from the hive for honey, pollen, propolis or water collect but they do not return to the hive.

Honeybees have magnetite crystal structures in body fat cells. These magnetite structures are the active components of the magneto-reception system. Thanks to these structures, honeybees can feel even slight changes in the magnetic field lines of the earth. These delicate structures are affected by the slightest magnetic pollution to occur and cause the honeybees to lose their direction. The bee dances that honeybees use to communicate with each other are distorted (Favre, 2011).

The electromagnetic field consists of electromagnetic waves. Electromagnetic waves consist of Electric Field and Magnetic Field components. These waves move at the speed of light.

Electromagnetic fields are physical fields produced by an Electro Magnetic field source. Electromagnetic waves are found in the continuous wavelength/frequency spectrum. The shorter the wavelength, the higher the frequency (Hernandez et al., 2010).

The Electro Magnetic Field is measured as the magnetic flux density and the unit is Tesla (T). The frequency of the electric magnetic fields is expressed in Hertz (Hz) (Vecchia et al., 2009).

Electro Magnetic field measurements can be influenced by different factors such as strength and distance of the source, the physical environment of the sites, the frequency of the radiation and possible modulation, reflection or polarization (Vecchia et al., 2009).

According to many studies, it has been reported that radio frequency and electromagnetic radiation (EMR) produce many misleading biological effects that disrupt the functions of all biological systems and all organisms (Blank and Goodman, 2009; Röösli et al., 2008; Schuz and Ahlbom., 2008)

The electromagnetic field can affect the immune system, working behavior and physiology of honeybees and ultimately cause them to disappear (Pattazhy, 2011). According to Sharma and Kumar (2010), a large amount of radiation also disturbs the bee’s ability to navigate and prevents them from returning to their hives. Honeybees are like a bioindicator of electromagnetic radiation because brain anatomies and learning regions are well known for associative learning abilities (Schwarzel and Muller, 2006). According to Pattazhy (2011), if the number of towers and mobile phones increases, honeybee may disappear within a decade.

According to the study, significant differences were found in returning to the hives of honeybees: 40 percent of the non- irradiated bees and 7.3 percent of the irradiated ones returned to their hives (Stefan et al., 2013). In this study, it was aimed to detect the effect of electromagnetic field intensity on the honeybees and waiting time of the bees in the area of the experiment.

## MATERIALS AND METHODS

The study was conducted on Caucasian honeybees (*Apis mellifera caucasica*). Caucasian bees are dark bees with gray hairs. They originated from the Caucasus mountains.

These bees are fairly gentle and have a longer language than other honeybee subspecies. They winter well in cold climates and raise strong colonies in the spring. Honey and Propolis production is more than other bee species and they are quite plundering but they are sensitive to *Nosema apis* and *Nosema ceranae.*

This study was carried out in 2017 at the Bayburt University Beekeeping Application and Research Station (40° 10□ 09□ N, 39° 50□ 53 26□ E). This Study was conducted in order to determine the effects of the electromagnetic field on honeybees’ time to locate food and their waiting time in the area.

In order to identify the numbers of bees that came to the Petri dishes for feeding, the experimental setup was placed at a distance of 100 m from the 50 caucasian hybrid bee colonies in the bee yard. Helmholtz coil equipment were placed in the rear of the bee yard, with a distance of 1.5m between them. In order to prevent chaos between the beehives and to make it easier to work, the back part of bee yard was preferred (Fig. 1).

**Figure 1.**
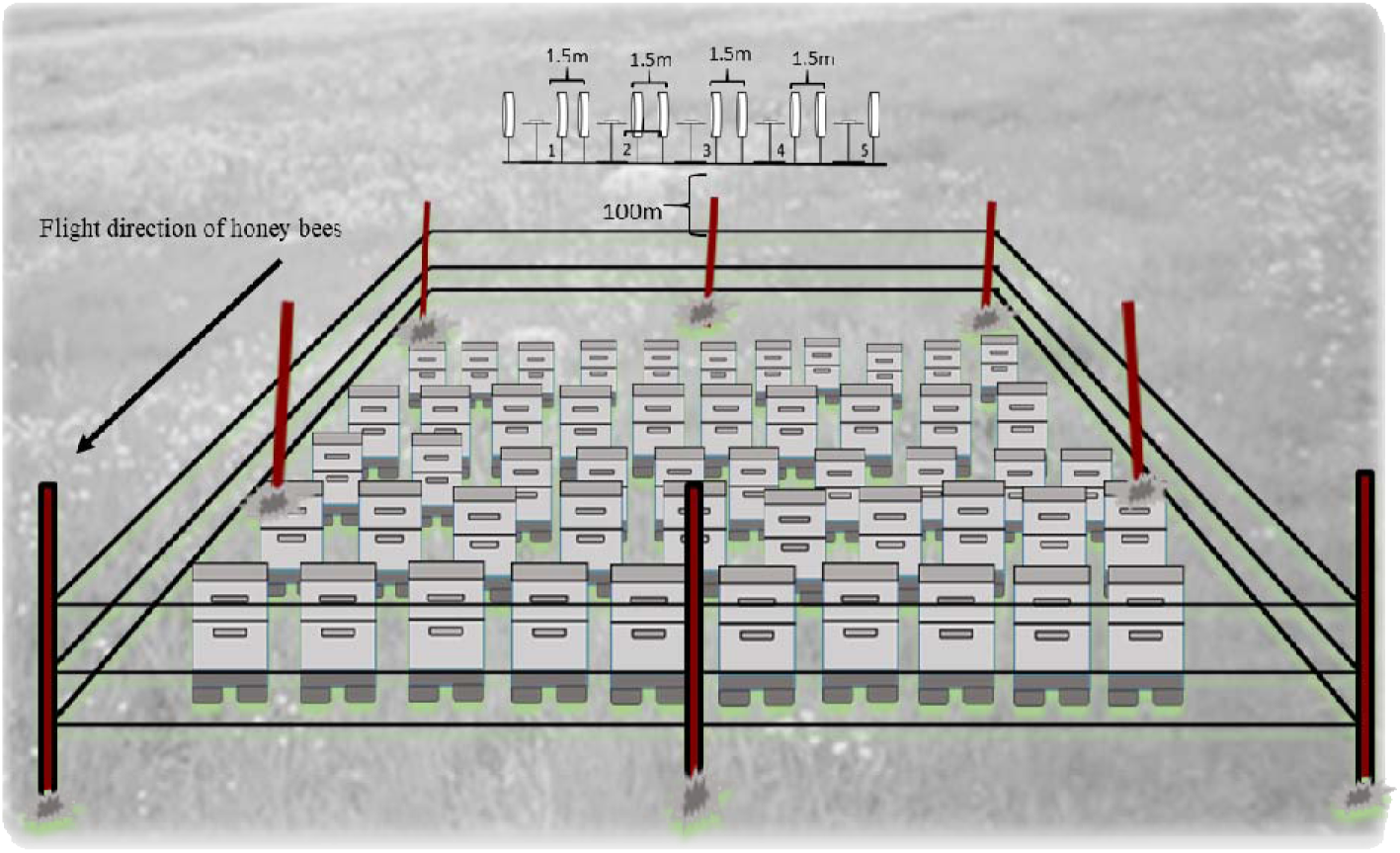
Positions of Helmholtz coil equipment according to apiary.

Helmholtz coil equipment was used to create electric and magnetic fields. Five Helmholtz bobbins and five different magnetic field levels were used in the study (Table: 1). The magnetic field strength produced by the Helmholtz coil equipment is adjusted by changing the voltage of the electricity applied to the coils. In this study, 50 Hz AC electricity was used.

**Table:1.**
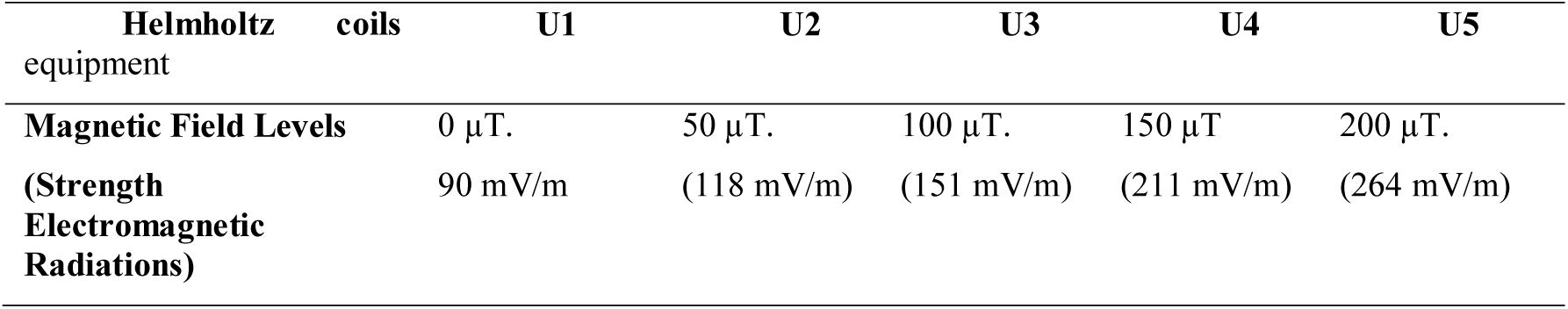
Application groups.

The Electromagnetic Field generated by the Helmholtz coil equipment was measured in terms of μT with the help of a TES Magnetic Field Meter model 1393. A diagram of the experimental setup is shown in Fig. 2. When the electromagnetic field is generated, the electric field also occurs at the same time. Both have an impact on living things. The strength of Electro Magnetic Radiation generated by Helmholts coils equipment is measured in terms of mV / m with the help of TES Electrosmog meter brand, model 593.

**Figure 2.**
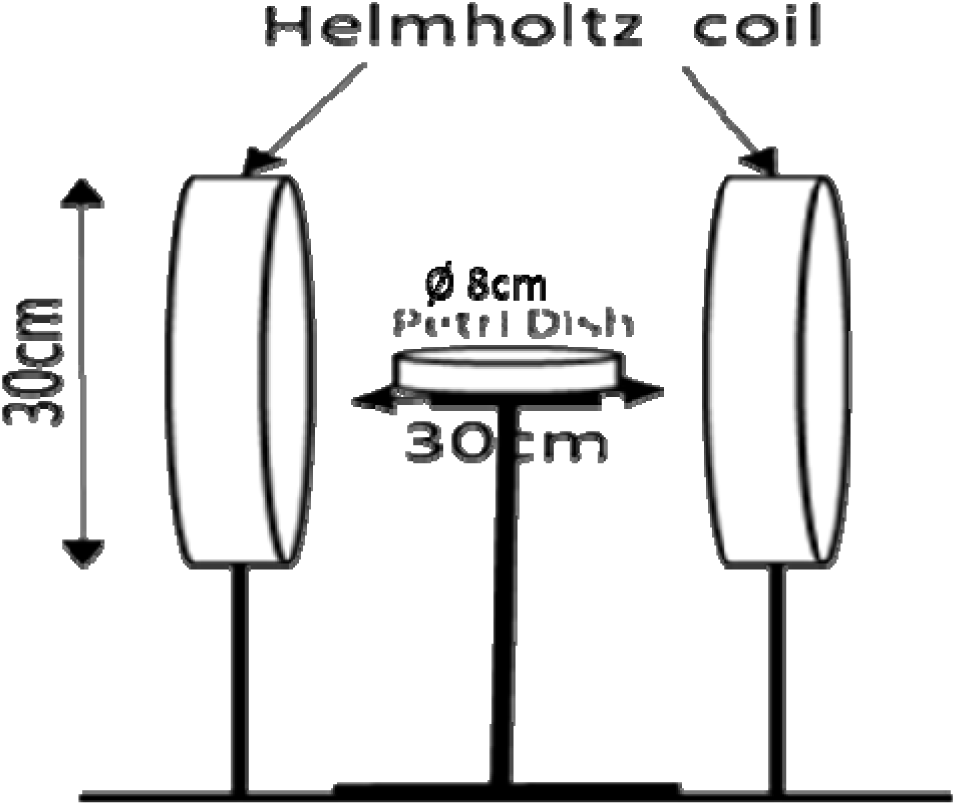
Representation of the Helmholtz coil equipment.

Petri dishes containing 25 cc 1: 1 syrup was placed in the center of Helmholtz bobbins and the experimental setup was prepared. The study began in the second week of June. Count down of honey bees were made between 14-16 o’clock. Because in the region where the study was conducted, the most intense nectar in this time range is carried. All of the helmotz coils equipment were energized at 14 o’clock at the same time and power was cut off at 16 o’clock.

The honeybees from the Petri dish were observed one by one, the period of time spent on the Petri dish was determined and recorded as the waiting period. This process was repeated 3 times with at least at15 day intervals. I took care to make the honeybees count down on rainyless and windless days. The juxtaposition of the Helmholtz coils equipment was done every time by draw lots.

The number of honeybees that came to feed in Petri dishes was detected, by counting with a minute interval. The counting process was continued until the syrup in the Petri dish was finished. All statistical analyses were performed using SPSS statistical software (IBM SPSS Statistics 22).

## RESULTS

As a first observation at the beginning of the study, honeybees began to circulate around petri dishes, but they did not alight on in Petri dishes. The first honeybee alight on Petri dish U1 (control (0 μT, 90 mV/m) after 5 minutes, followed by U2 (50 μT, 118 mV/m), U3 (100 μT, 151 mV/m), U4 (150 μT, 211 mV/m), and U5 (200 μT, 264 mV/m). The most visited, application was U1 (0 μT, 90mV/m) (mean 21.07±17.89 bees) and the least visited application was U5 (200 μT, 264 mV/m) (mean 10.82±11, 77 bees) (Table 2). Honeybees have passed intensely on Petri dishes with a magnetic field at the top after finishing the feed in the Petri dish. As the magnetic field intensity increases, the demand for honeybees decreases and reluctance is seen (Table 1). Although the bees placed in the U1 (0 μT, 90 mV/m) Petri dish where no magnetic field was present stayed here for longer (mean 37.88 s) (Table 1), the bees placed in U5 (200 μT, 264 mV/m) Petri dishes with high magnetic field abandoned Petri dishes in much shorter time (mean 12.61 Sec).

**Table 2.**
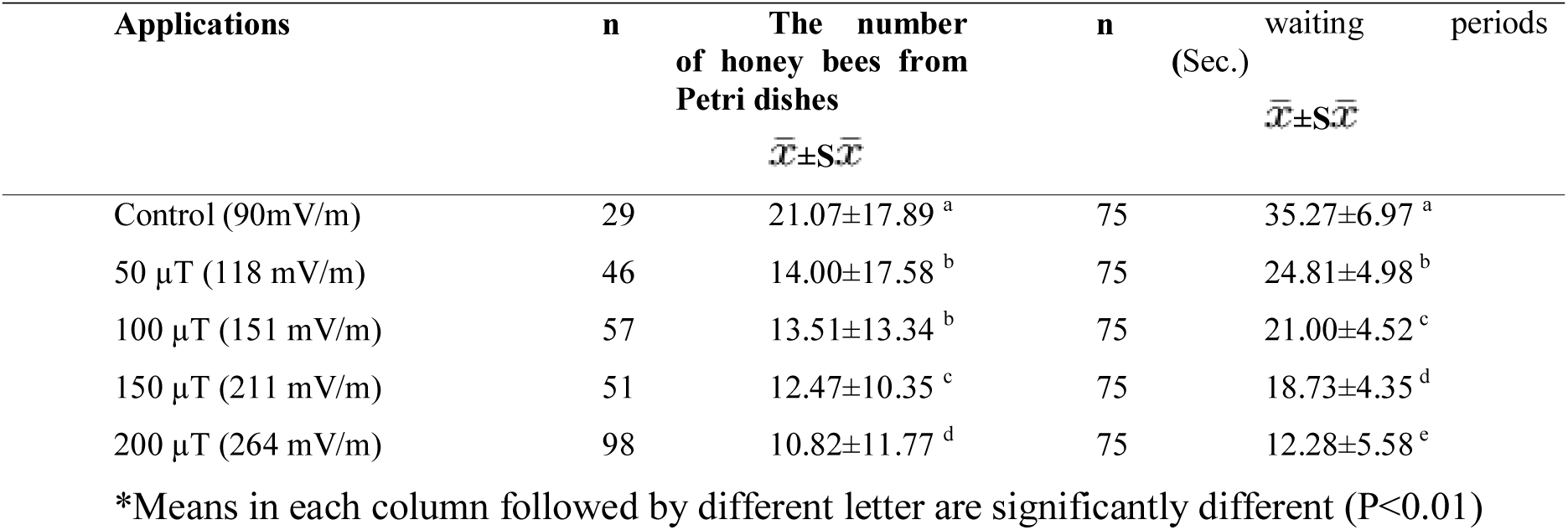
Average values of the number of honeybees from Petri dishes and waiting periods in Petri dishes.

From the multiple comparison tests conducted, it was found that the application groups were located in different groups (Table 2).

Analyzes of variance were made for bee numbers in Petri dishes (Table 3) and for the time they spent in Petri dishes of honey bees (Table 4).

**Table 3:** Results of variance analysis on honeybee numbers in Petri dishes.

| Variation Sources | df | Mean Square | F | Sig. |
| --- | --- | --- | --- | --- |
| Minutes | 32 | 884.793 | 3539.171 | .000 |
| Iteration | 2 | 34.722 | 138.888 | .007 |
| Applications | 4 | 3485.440 | 13941.762 | .000 |
| Minutes * Iteration | 64 | 11.250 | 45.001 | .022 |
| Minutes * Applications | 70 | 167.961 | 671.843 | .001 |
| Iteration * Applications | 8 | 75.340 | 301.361 | .003 |
| minutes * Iteration * Applications | 98 | 10.838 | 43.350 | .023 |
| Error | 2 | 0.250 |  |  |

**Table 4:** Results of analysis of variance applied to the period in which honeybees spend their Petri dishes.

| Variation Resources | f | Mean Square | F | Sig. |
| --- | --- | --- | --- | --- |
| Applications |  | 5422.357 | 222.250 | .000 |
| Iteration |  | 3.523 | .144 | .866 |
| Iteration* Applications |  | 7.959 | .326 | .956 |
| Error | 60 | 24.398 |  |  |

According to these results, while the highest number of bees and the waiting time were 0 μT magnetic fields applied Petri dishes, the number of bees that alight on and the duration of stay was the lowest of 200 μT magnetic fields applied Petri dish (Table 2).

## DISCUSSION

Different studies from different regions of the world have reported the negative effect of EMF emitted from cell phone towers, high voltage wires and various electronic devices on honey bees with regard to strength, navigation, behavior, honey store, pollen store and brood area, etc. (Harst et al., 2006; Stefan et al., 2013; Sharma and Kumar, 2010; Pereira-Bomfim et al. 2015).

However, some other researchers have reported that EMF has no effect on honeybees (Mixson et al., 2009; Blacquiere and Hoofwijk, 2010).

According to a study conducted by Mall and Kumar, the bee colonies were not affected by EMF but reported that they could damage honeybees in the long term (Mall and Kumar, 2014).

Studies on the effects of electromagnetic fields on honey bees have shown that initiation of foraging, cessation of foraging and number of incoming foragers are negatively affected (Harst et al., 2006; Kimmel et al., 2007; Stefan et al., 2013; Sharma and Kumar, 2010; Pattazhy, 2011; Darney et al., 2016; Taye et al. 2017) the number of outgoing foragers (Valberg, 2010; Sharma and Kumar, 2010), the successful return of marked feeders (Harst et al., 2006; Stefan et al., 2013)

On the contrary, a few researchers have argued that the EMF does not have a negative effect on honeybees (Mixson et al., 2009; Blacquiere and Hoofwijk, 2010; Singh, 2014).

Considering these studies, a study entitled “The Effect of Electromagnetic Field (EMF) on Nutritional Behavior of Honey Bees (*Apis mellifera* L.)” was conducted at Bayburt during 2017.

The study on the number of incoming bees increased from U1 (control 0 μT, 90 mV/m, 10.82±11.77 bees) to U5 (200 μT, 264 mV/m, 21.07±17.89 bees). It was observed that when the EMF or electric field intensity increases the number of bees that arrived in the Petri dishes and the waiting time of them decreases (Table 2).

That is, the EMF or electric field intensity increases, the number of bees from petri dishes and the waiting time of Petri dishes decreases.

## Conclusion

The present results showed that honeybees are sensitive to the modification of EMF or electric field intensity.

Recently, Valkova and Vacha (2012) discussed the possibility of using honeybees for both magnetic nanoparticles and the magnetic field of the earth to detect the geomagnetic field.

In conclusion, Honeybees have been observed for the first time under the influence of electric and electromagnetic fields. Firstly, honeybees have been added to the list of animals that have been studied on magnetoreception and electroreception.

It can be deduced from our results that areas where the electromagnetic field is dense will be less visited by bees, resulting in the fact that plants and fruit trees in these regions will not be sufficiently pollinated. This will cause a decrease in the quality of fruits and other plant products.

The development of technology increased by electromagnetic pollution will effect honeybees and crop production negatively. The apiaries should be installed away from high-voltage lines, base stations, industrial zones and residential areas in order to reduce the negative impact of the electromagnetic field or the electric field on honeybees.

## Acknowledgments

### Competing interests

There is no conflict of interest.

### Funding

The authors thank the Bayburt University for their financial support

